# Better ears with eyes open: effects of multisensory stimulation with nonconscious visual stimuli on auditory learning

**DOI:** 10.1101/519900

**Authors:** Milton A. V. Ávila, Rafael N. Ruggiero, João P. Leite, Lezio S. Bueno-Junior, Cristina M. Del-Ben

## Abstract

Audiovisual integration may improve unisensory perceptual performance and learning. Interestingly, this integration may occur even when one of the sensory modalities is not conscious to the subject, e.g., semantic auditory information may impact nonconscious visual perception. Studies have shown that the flow of nonconscious visual information is mostly restricted to early cortical processing, without reaching higher-order areas such as the parieto-frontal network. Thus, because multisensory cortical interactions may already occur in early stages of processing, we hypothesized that nonconscious visual stimulation might facilitate auditory pitch learning. In this study we used a pitch learning paradigm, in which individuals had to identify six pitches in a scale with constant intervals of 50 cents. Subjects were assigned to one of three training groups: the test group (Auditory + congruent unconscious visual, AV), and two control groups (Auditory only, A, and Auditory + incongruent unconscious visual, AVi). Auditory-only tests were done before and after training in all groups. Electroencephalography (EEG) was recorded throughout the experiment. Results show that the test group (AV, with congruent nonconscious visual stimuli) performed better during the training, and showed a greater improvement from pre-to post-test. Control groups did not differ from one another. Changes in the AV group were mainly due to performances in the first and last pitches of the scale. We also observed consistent EEG patterns associated with this performance improvement in the AV group, especially maintenance of higher theta-band power after training in central and temporal areas, and stronger theta-band synchrony between visual and auditory cortices. Therefore, we show that nonconscious multisensory interactions are powerful enough to boost auditory perceptual learning, and that increased functional connectivity between early visual and auditory cortices after training might play a role in this effect. Moreover, we provide a methodological contribution for future studies on auditory perceptual learning, particularly those applied to relative and absolute pitch training.

## 1. Introduction

Musicians have different brains compared to non-musicians. In fact, auditory processing by musicians has been linked to distinctive electrophysiological markers, particularly changes to event-related potentials (ERP) (e.g. Fujioka et al., 2004; Menning et al. 2000; Seppanen et al., 2012). In addition, magnetic resonance imaging (MRI) studies demonstrate that music recruits a brainwide network encompassing not only auditory, but also motor and associative areas (e.g. Gaser & Schlaug, 2003; Zatorre et al., 2007; Schlaug et al., 2010; Schlaug, 2015). This is actually not surprising, since music is a multisensory phenomenon (Zimmerman & Lahav, 2012). Vision, for instance, has been demonstrated to influence appreciation and judgement of musical tones (Schütz, 2007), as well as musical performance (Tsay, 2013).

Musical appreciation and performance is just one instance of multisensory integration, which in a broader sense is present in our everyday lives. In fact, multisensory integration reflects the ability of our brains to bond different, but congruent, sensory inputs into a common percept. A necessary condition for integration is the temporal alignment of sensory streams (Jones & Jarick, 2006; Shore et al., 2006; van Wassenhove et al., 2007). Also fundamental is the correspondence of sensory features, e.g. size and timbre, size and pitch, and the particularly well studied correspondence between spatial elevation and pitch (Melara & O’Brien, 1987; Ben-Artzi & Marks, 1995; Patching & Quinlan, 2002; Maeda et al., 2004; Widmann et al., 2004; Evans and Treisman, 2010; Spence, 2011; Salgado-Montejo et al., 2016). Interestingly, this ability may still be true when one of these features is nonconscious. In fact, studies have demonstrated that the processing of a visual stimulus that is not consciously perceived may be influenced by an auditory stream (Chen & Spence, 2010, 2011; Ngo & Spence, 2010; Palmer & Ramsey, 2012; Alsius & Munhall, 2013).

There has been a great effort toward understanding the neural mechanisms of conscious visual perception, but a consensus is far from being achieved (Crick & Koch, 2003; Lamme, 2006; Cohen & Dennet, 2011; Fahrenfort & Lamme, 2012). Studies suggest that this phenomenon relies on subcortical structures (e.g. Paus, 2000), primary cortical areas (Zeki, 2003; Lamme 2003), or even association areas such as parieto-frontal network (Dehaene & Naccache, 2001; Rees et al., 2002). In this context, Dehaene et al. (2006) have proposed a model of visual consciousness with three stages of processing: subliminal, pre-conscious, and conscious. The main difference between them relies on both the strength of the activation of visual cortices and the likelihood with which this activation is spread to higher order association areas. Basically, the model proposes that in nonconscious processing cortical activation does not reach parieto-frontal networks, and instead remains restricted to early visual and occipito-temporal cortices (Dehaene et al., 2006).

Multisensory integration is known to involve heteromodal areas in association cortices, such as the Superior Temporal Sulcus (Beauchamp et al., 2004; Powers et al., 2012), the Intraparietal Sulcus (Grefkes & Fink, 2005), and frontal areas (Calvert, 2001). However, there is growing evidence that this integration also encompasses cortical areas traditionally regarded as unisensory (Ghazanfar & Schroeder, 2006; Liang et al., 2013, Murray et al., 2016). Evidences also point to the communication between primary cortical areas in multisensory events (Schroeder et al., 2004; Cappe & Barone, 2005; Foxe & Schroeder, 2005; Ghazanfar & Schroeder, 2006; Lewis & Noppeney, 2010), which is supported by anatomical connectivity studies in other animals (Clarke & Innocenti, 1990; Falchier et al., 2002, 2009; Rockland & Ojima, 2003; Cappe & Barone, 2005; Clemo et al., 2008; Laramée et al., 2011; Ibrahim et al., 2016). These connections are more robust in areas that process peripheral visual stimuli (Falchier et al., 2002; Rockland & Ojima, 2003), and may also involve subcortical structures and feedback mechanisms (Tyll, 2011; Proulx et al., 2012, for review). Noteworthy, with this growing interest on multisensory integration, new techniques and procedures for perceptual learning are being developed.

Most studies have focused on how visual perceptual learning is influenced by other modalities (e.g. Seitz et al., 2006; Kim et al., 2010; King, 2009; Powers et al., 2012, Zilber et al., 2014). These studies demonstrate that multisensory training is more efficient than unisensory training in improving perceptual performance. In addition, there are evidences that multisensory training induces neural plasticity in both unisensory and heteromodal areas (Zilber et al, 2014). This seems also true for musical training: Lappe et al. (2008) report that ERP mismatch negativity (MMN, a neurophysiological marker of neural plasticity) is stronger after multimodal (sensorimotor + auditory) than unimodal training (auditory only). Also, Paraskevopoulos et al. (2012) show MMN alterations in multisensory areas after multimodal training (Paraskevopoulos et al., 2012).

Therefore, considering that: i) even a nonconscious visual event may elicit activation in early sensory cortices, ii) multisensory integration may occur since early stages of processing, and iii) nonconscious perceptual learning has been demonstrated in the literature (e.g. Watanabe et al., 2001), we hypothesized that nonconscious visual stimulation might improve auditory pitch learning. More specifically, we aimed at responding two main questions: 1) Would a pitch learning paradigm benefit from a multisensory training protocol? 2) If so, would early “unisensory” cortices be involved in training-induced alterations in multisensory networks?

## 2. Materials and Methods

### 2.1. Participants

After providing written informed consent, 34 subjects (19 males) with ages between 18 and 40 years (all of which right-handed, non-musicians, with normal or corrected-to-normal vision, and no prior neurological disorders), participated in the experiment. One participant, who reported clear visual perception during training was excluded from the experiment. The protocol was approved by the Research Ethics Committee, General Hospital of the Ribeirao Preto Medical School, University of Sao Paulo (CAAE 34853614.0.0000.5440).

To control for the effect of musical experience, only subjects with no prior formal musical education were recruited. Before the experiment, they responded to a questionnaire in order to estimate the Musical Sophistication Index (MSI) (Mullensiefen *et al.,* 2014). Both musical perception and musical training indices were took into account when balancing the groups.

Participants were pseudorandomly assigned to three groups: Auditory-only (A), trained only with auditory information; Audiovisual (AV), trained with auditory + nonconscious visual information; and Audiovisual Incongruent (AVi), trained with auditory + nonconscious and spatially incongruent visual information. Table 1 describes groups according to mean age and musical indices. No statistically significant differences were found.

**Table 1.** Group description

|  | **n** | **age** | **musical perception**<br>**(1 to 7)** | **musical training**<br>**(1 to 7)** |
| --- | --- | --- | --- | --- |
| **A** | 11 | 27.7 ± 3.41 | 4.67 ± 0.75 | 2.41 ± 1.26 |
| **AV** | 11 | 25.8 ± 4.5 | 4.27 ± 1.14 | 1.70 ± 0.95 |
| **AVi** | 11 | 26.0 ± 6.6 | 4.20 ± 1.13 | 1.93 ± 1.08 |

### 2.2. Apparatus

Stimuli were generated using a custom Python 2.7 script running on a desktop with an i3 processor, OS Windows XP, NVidia GTX960 graphics card. Stimuli were presented on virtual reality goggles (Oculus Rift DK1) at 60 Hz refresh rate (i.e., frames per second), and resolution of 1280 x 800 pixels. We used these virtual reality goggles to assure visual stimuli were presented at a fixed visual angle while allowing participants to sit comfortably. Screen luminance was 92 cd/m^2^. Auditory stimuli were presented on over-the-ear headphones (Audio-Technica model ATH-M10).

### 2.3 Experimental design

The experiment comprised two daily sessions, with approximate duration of 45 min on Day 1, and 55 min on Day 2. Day 1 aimed at identifying the visual detection thresholds of each individual (Figure 1). Day 2 was the experiment proper, consisting of a pitch learning paradigm based on a 6-alternative forced choice task (6AFC). Electroencephalography (EEG) was recorded throughout Day 2 (Figure 1).

**Figure 1.**
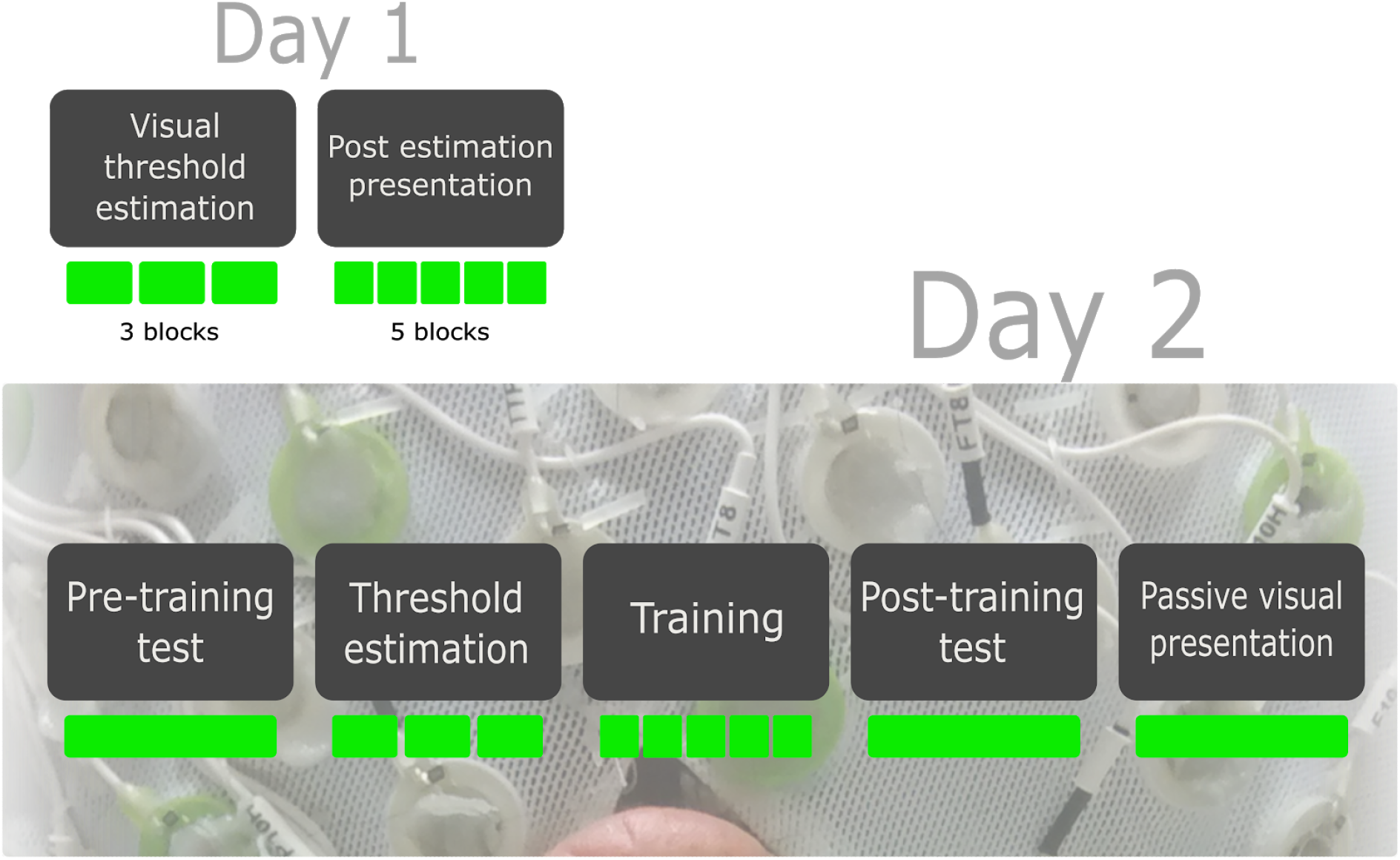
Experimental timelines. Day 1 was divided in two phases. First phase: three blocks of staircase procedure to estimate detection thresholds. Second, five blocks of visual stimulation. Day 2 was divided in five phases, all of them with EEG recording. First phase: pre-training test. Second, a staircase procedure like on Day 1. Third, five blocks of pitch training. Fourth, Post-training test. Fifth, visual stimuli presented during 3 minutes, with no motor responses.

Each visual stimulus consisted of a 1-pixel flash, corresponding to 0.13° of visual angle, presented for two refresh cycles (i.e., approximately 33 ms duration, given the 60 Hz frame rate). Each flash was presented in six possible locations on the screen, as illustrated in Figure 2. A non-informative auditory beep (1000 Hz, 33 ms, 70 dB, 5 ms of onset and offset ramps) was paired with each visual stimulus. Threshold estimation was done using a staircase procedure (Garcia-Perez, 1998) (Figure 3A).

**Figure 2.**
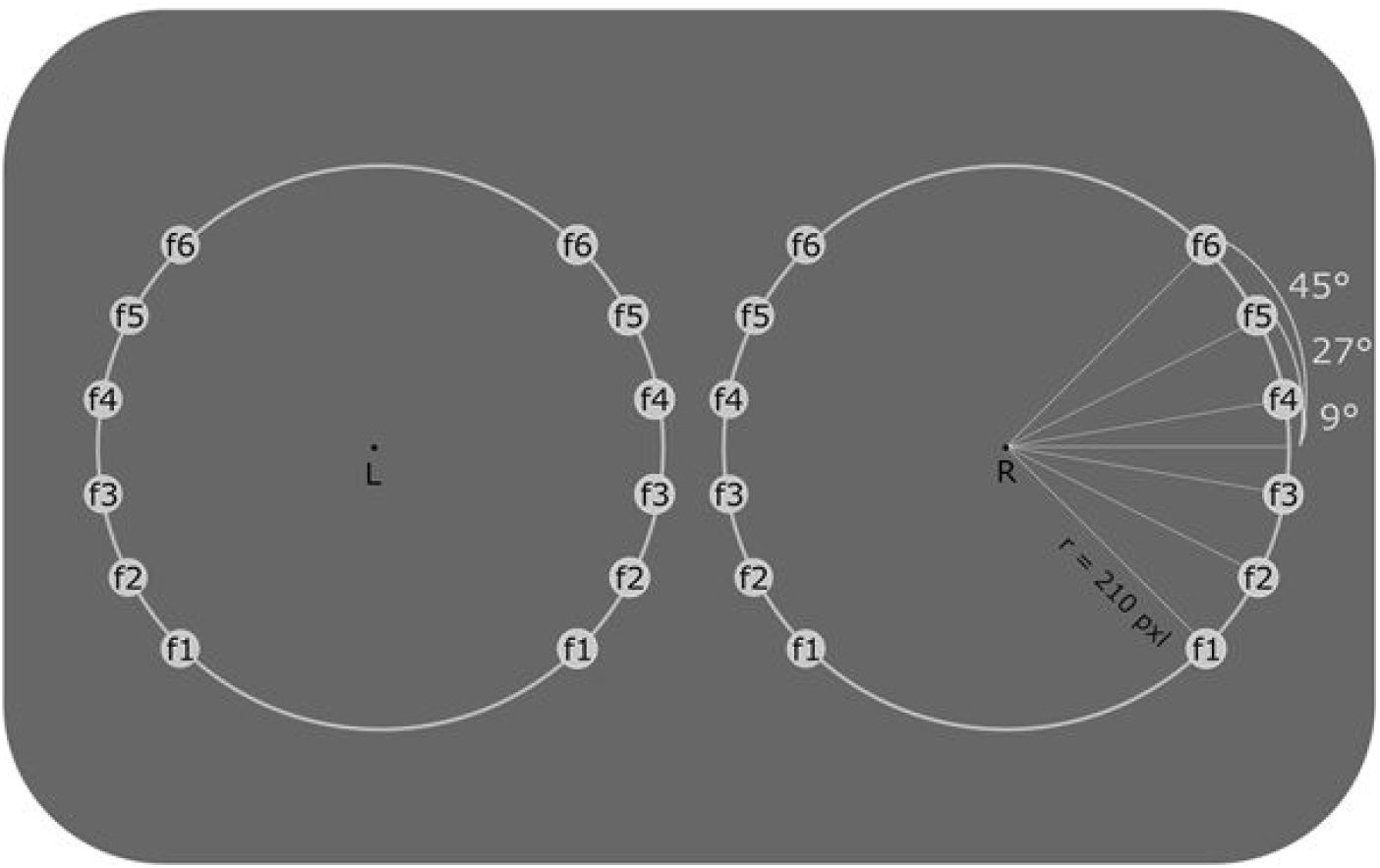
Visual stimuli. The same stimuli were presented to the left and right eyes on a dark gray background. Each flash corresponded to an auditory frequency temporally aligned with the tone. Flashes 1 to 6 were equally distributed in 90° of a circumference with 210 pixels of radius (28° visual angle). Such radius had been determined in a pilot study with 33 subjects. The illustrated circumference was not presented. Distance from L and R was established according to the interpupillary distance of each subject.

After the staircase procedure, subjects were submitted to five blocks of visual stimulation (Figure 3). Stimuli were presented in seven different contrasts: 0% (the threshold contrast), −10%, −20%, −30%, −40%, −50%, or-100% (the “catch” trials). Subjects were asked to respond whether the stimulus was present or not in each trial. After calculating the d’ of each pair (Green & Swets, 1966), we ran 100 thousand simulations of an observer with zero sensitivity doing the same task. As a result we obtained a distribution of d’ to determine which contrast levels yielded a performance significantly greater than expected from blind guessing. The highest contrast with a d’ lower than that corresponding to 0.01 of probability was taken as reference for the contrast to be presented during training. To calculate contrasts during training, the reference was multiplied by 0.75. At the end of Day 1, participants were asked to answer the MSI questionnaire.

**Figure 3.**
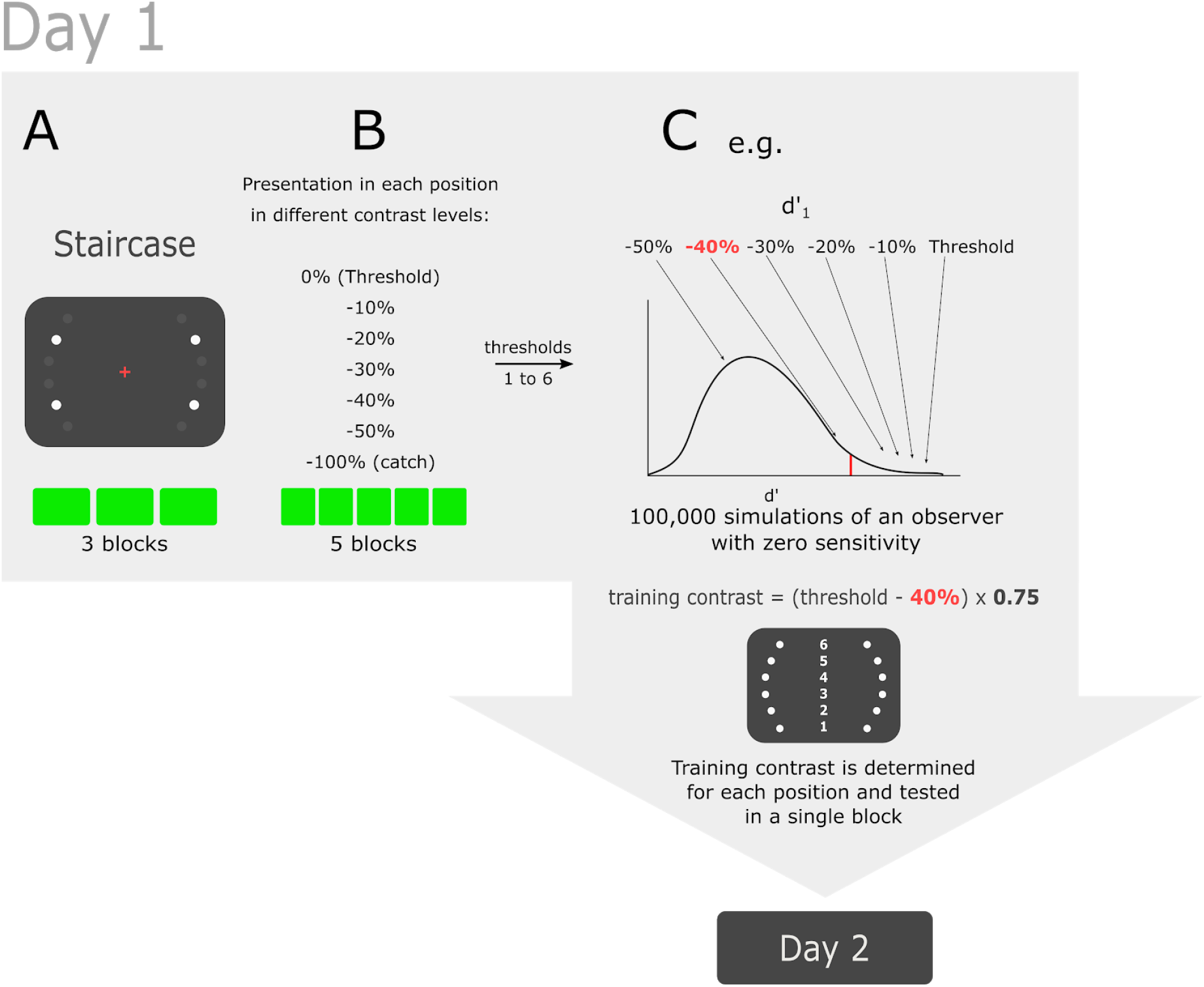
Day 1 protocol. **A.** Three staircase blocks for threshold estimation were run. **B.** Presentation of all stimuli in different contrast levels: 0% (i.e., threshold), −10%, −20%, −30%, −40%, −50%, −100% (i.e., catch trials). Stimuli were presented in all six positions randomly. For each contrast level and each position, the d’ was determined according to the Signal Detection Theory (Green & Swets, 1966). **C.** After obtaining a distribution of 100 thousand simulated observations, we estimated the probability of each d’ to occur by blind guessing. The reference for training contrast was the highest contrast level required for a d’ for which the probability was lower than 0.01. The training contrast was determined by multiplying the reference contrast by 0.75.

Before beginning Day 2, the subject was prepared for EEG acquisition, received written instructions, and was given the opportunity to ask questions about the experiment. Instructions were repeated on the screen along the experiment, prior to beginning the next phase. Participants sat comfortably, and could rest between blocks.

Day 2 is depicted in Figure 4, and is described in detail in the next subsections.

**Figure 4.**
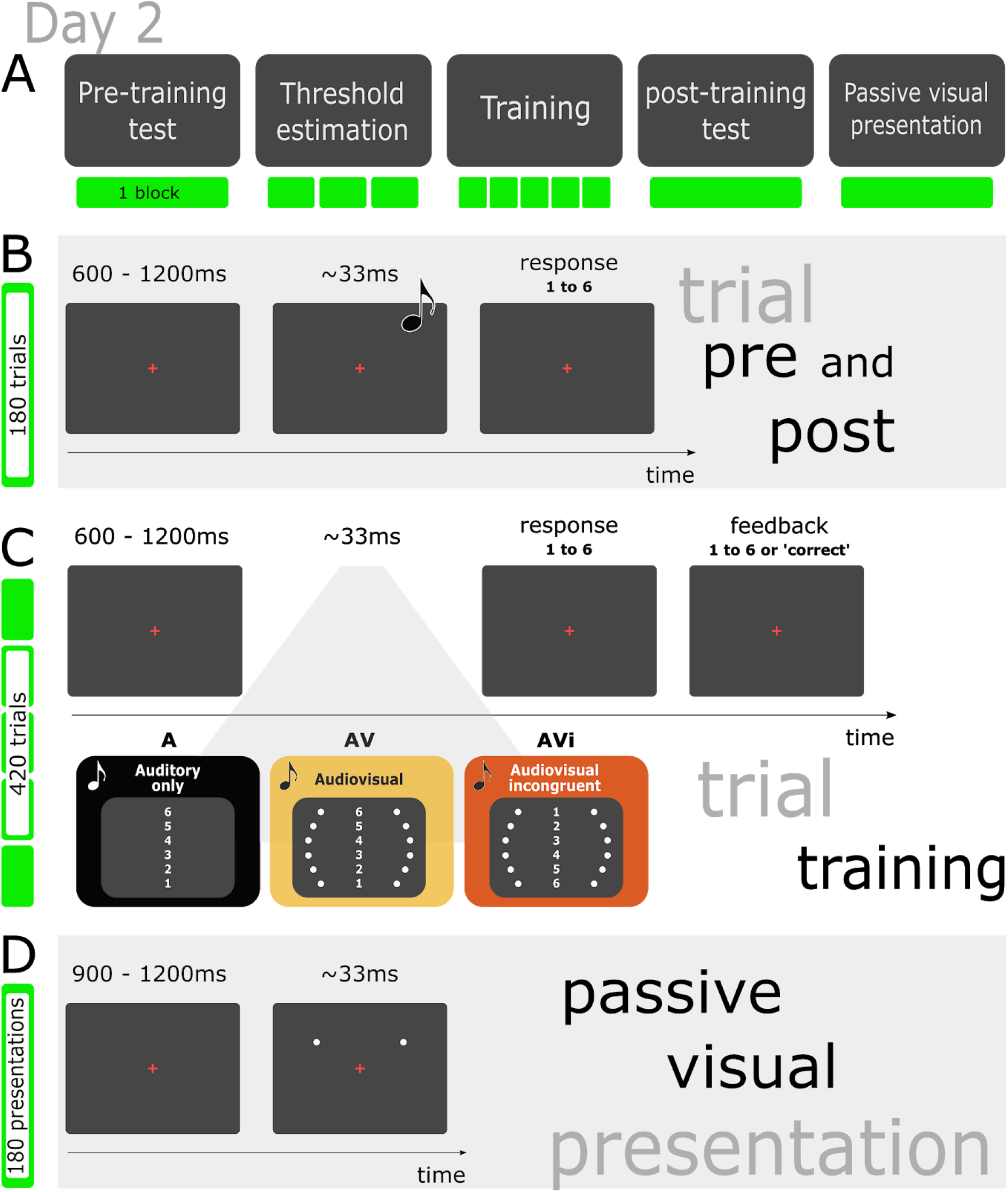
Day 2 protocol. **A.** Overview of the timeline. **B** Pre- and post-training tests. Each trial started with the presentation of a fixation cross for 600-1200 ms, followed by an auditory beep of 33 ms. Participants had to inform as quickly as possible (in a 6AFC schedule) which frequency was played (from 1 to 6). Each test consisted of 180 trials. **C.** Training. Similar to pre- and post-tests, but with visual stimuli paired with auditory stimuli, depending on the group of subjects. Group A: auditory stimuli alone. Group AV: auditory stimuli with spatially congruent (and nonconscious) visual stimuli. Group AVi: auditory stimuli with spatially incongruent (and nonconscious) visual stimuli. The numbers indicate what frequency (1 to 6) was paired with what pair of nonconscious flashes. A feedback was given to the subject after each response, displaying either the word “correct” or the actual frequency number if the response was incorrect. A total of 420 trials were presented across 5 blocks. **D.** Passive visual presentation. Similar to pre-and post-tests, but without demanding responses.

#### 2.3.1. Pre- and post-training tests

The pre- and post-training tests consisted of 180 trials with 30 presentations of each auditory frequency. Each trial started with a fixation cross in the center of the screen for 600-1200 ms. The auditory stimulus was then presented for 33 ms (70 dB). The frequency of each auditory stimulus was randomly selected among six different frequencies (Table 2). After the auditory stimulus, the participant had to inform as quickly as possible (in a 6AFC schedule) which frequency had been presented (Figure 4B). All responses were collected on a masked keyboard, so that only the desired keys were available. Responses for tone 1 to 3 were given with the left hand, and 4 to 6 with the right hand. Reaction times (RT) consisted of the time elapsed between stimulus and response. Before the test began, the participant heard the tone and saw the corresponding frequency three times each. Then s/he performed a familiarization task, with 4 presentations of each frequency followed by the response.

**Table 2.** Frequencies and their corresponding numbers

| 1 | 2 | 3 | 4 | 5 | 6 |
| --- | --- | --- | --- | --- | --- |
| 1192.1Hz | 1227.0Hz | 1263.0Hz | 1300.0Hz | 1338.1Hz | 1377.3Hz |

Auditory stimuli were computationally produced as sine waves. Frequencies and stimulus duration were designed to be barely distinguishable by normal-hearing non-musicians (Plack et al., 2005), thus making it difficult for subjects to achieve ceiling performance. All pitch intervals were a quarter tone, or 50 cents, and none of the frequencies we used belong to a Western tempered scale.

At the end of each phase, participants were asked, in a scale from zero to 10, how confident they were about their performance, which yielded us confidence ratings.

#### 2.3.5. Threshold estimation and confirmation

After the pre-training phase, thresholds were estimated as on Day 1 to check how consistent were contrast levels between days. A <10% variation from Day 1 to Day 2 was tolerated. Otherwise, the participant was placed in Group A, i.e., with an auditory-only training. Importantly, this was the case of only one subject.

#### 2.3.6. Training phase

The training consisted of 420 randomly sequenced trials, 70 for each frequency, divided in five blocks. It was during the training that participants were divided into the three experimental groups. Group A was trained solely with auditory stimuli. Group AV, in turn, was trained with spatially congruent visual stimuli. By congruence we mean that sound frequency and visual elevation had the same direction, e.g., frequencies 1 and 6 were paired with lowest and highest positions, respectively. Conversely, Group AVi was trained with spatially incongruent visual stimuli. In other words, sound frequency and visual elevation directions were opposite to each other, e.g., frequencies 1 and 6 were paired with the highest and lowest positions, respectively (Figure 4C). With such design we approached the hypothesis that incongruence between two sensory modalities impairs multisensory integration at the perceptual level.

On Day 1, participants were informed that the experiment had two main focuses: vision on Day 1, and audition on Day 2. This was an attempt to direct their attention to auditory information on Day 2, thus favoring visual stimuli to be nonconscious (Dehaene et al., 2006). To reinforce that even further, before Day 2 session began, participants were informed that an auditory learning test was about to happen, and so the only important information was the auditory tones they had to identify, except for the staircase phase. Finally, after training, participants were asked whether they saw any flash. If so, they were asked to estimate such perception on a percentage scale from 0-10% to 90%-100%. Any subject reporting that they clearly saw visual flashes was supposed to be excluded from analysis. This indeed happened in one individual, who reported 70% - 80%. Other three individuals reported 0 - 10% and, interestingly, two of them were in group A, i.e., they actually had not been visually stimulated. We therefore decided to keep these three participants in the analysis. None of the other participants reported having seen any flashes.

#### 2.3.7. Passive visual presentation

In this final phase of Day 2, participants were exposed to the same visual stimuli presented during the training, but without auditory stimulation. No responses were required, so their task was to simply focus their gaze at the fixation cross for approximately 3 minutes. The aim was to collect a basal cortical response evoked by the visual stimuli alone, making it possible to disentangle auditory and visual cortical responses collected in previous phases.

### 2.4. Behavioral analysis

Performances were calculated based on the distance the response was from the tone presented in each trial. The average of these distances resulted in Mean Absolute Deviations (MAD) (Bermudez et al., 2008). MAD, was calculated according to Equation A, where ***x*** denotes the difference between stimulus and response, and ***n*** denotes the number of presentations.

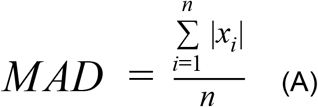

Performances were calculated for pre- and post-training tests using the MAD. The effect of training was the ratio between pre- and post-training tests, given in percentage of performance improvement. Group comparisons were made with the non-parametric Kruskal-Wallis test, since distributions were not normal according to the Shapiro-Wilk test. For *post hoc* analyses we used the Wilcoxon-Mann-Whitney test. We also calculated Cohen’s d for effect sizes. Performance during training was compared between groups by applying the same statistical analyses.

For a deeper investigation on behavioral performances, we calculated MAD for each auditory frequency separately. Distributions were then fit with a quadratic function. Of note, frequencies 1 and 6 were observed to induce the clearest behavioral improvements. Hence, we decided to sort auditory stimuli as outer (1 and 6) and inner frequencies (2 to 5). The effect of training was then calculated for each category of pitches and compared between groups of subjects.

Confidence ratings obtained after two phases, pre- and post-training, were compared between groups, as well as the pre/post-training ratio. RT were also compared between groups of subjects in each phase. Trials with RT below 150 ms and above 2300 ms (i.e., with potentially spurious responses) were discarded. Because we found differences in RT, we tested multiple correlations among RT, and the effects of training, MSI variables, and confidence ratings.

### 2.5. EEG acquisition

We acquired EEG using a 128-channel cap (Brain AMP, Brain Products, Guilching, Alemanha), with Ag/AgCl electrodes, internal resistance of 5 kΩ, and FCz as the reference electrode. To reduce impedance we used ABRALYT HiCl gel during montage aiming at an impedance below 20 kΩ for all electrodes. Signals were digitized at 5 kHz.

### 2.6. Preprocessing

For both preprocessing and analysis, we combined eeglab (Delorme and Makeig, 2004), chronux (Mitra and Bokil, 2007), and custom scripts in Matlab. Signals were initially downsampled to 500 Hz, and band-pass filtered (0.5-50 Hz) using the digital FIR filter. Noisy channels were rejected through visual inspection. Good channels were re-referenced based on the average of all electrodes, and rejected channels were then interpolated. Artifacts were rejected using both visual inspection and independent component analysis (ICA). Trials with artifacts surviving this scrutiny were manually rejected. On average, 24.6 trials per subject were rejected. Accepted trials were epoched in 4400 ms stimulus-locked sweeps: 1900 ms before, and 2500 ms after stimulus onsets. These sweeps included: (1) the actual perievent window described in the Results section (400 ms before, 1000 ms after stimulus onsets), plus (2) margins for time-frequency analysis (1500 ms before, 1500 ms after such perievent window).

### 2.7. EEG analysis

#### 2.7.1. Time-frequency analysis

Time-frequency decomposition was performed though Morlet wavelet convolutions. Power was calculated both from the raw EEG and after applying the Laplacian filter. Fifty frequencies between 1 and 50 Hz were extracted from the time series, using logarithmically spaced cycles between 4 (1 Hz) and 12 (50 Hz).

#### 2.7.2. Connectivity

For connectivity analysis, all EEG epochs were converted to Surface Laplacian (also known as Current Source Density), as proposed by Perrin et al. (1989). The three parameters for this conversion were smoothing (lambda), Lagrange order (number of iterations when computing the Legendre polynomial), and spherical spline order (m), respectively set to 10^-5^, 10, e 4 (Cohen, 2015).

To estimate the Interchannel Phase Synchrony (ICPS), we obtained the difference of phase angles between two electrodes by calculating their cross-spectrum via the multi-taper method. In this method, different orthogonal tapers (slepian sequences) are applied to the signal, and the FFT (fast fourier transform) is calculated for each signal using a moving window, resulting in values with time and frequency variation. Assuming *z_k_*(*f*) and *z_j_*(*f*) as the results of the complex Fourier transform of the temporal series *z*’_*k*_(*t*) e *z*’_*j*_(*t*) from channels 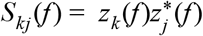, where * denotes the complex conjugate. Thus, ICPS was given as in equation B:

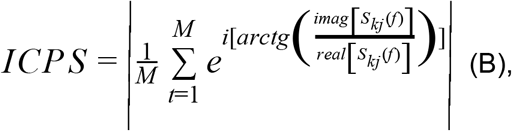

where M denotes the number of temporal windows used to calculate *S_kj_* (*f*), which assumes a different value for each temporal window. We used 0.5 s windows with 50% overlap, resulting in 25 M values (bandwidth product of 3 and 5 tapers).

#### 2.7.3. Statistics

For power values and ICPS we averaged trials and normalized to a decibel (dB), as in equation C:

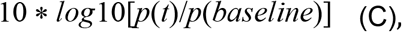

allowing direct comparison of effects on frequency bands. We then averaged across the subjects of each group and calculated the difference between pre- and post-training tests. In order to perform group comparisons, we calculated the power and ICPS in the theta band (3-8 Hz). To test data normality we used the Kolmogorov-Smirnov test, and for group comparisons we used one-way ANOVA (when three groups were being compared) and unpaired Student’s t-test (for two groups).

## 3. Results

### 3.1. Behavior

To verify whether pitch perceptual learning benefits from multisensory training, we analyzed performances during three phases: pre-training, training, and post-training. The lower was the MAD, the better was the performance. Thus, it can be seen from Figure 5A that all groups improved their performance after training. No significant effects during training were associated with the main graph of Figure 5A, which depicts performances across phases and training blocks (H = 3.65, p = 0.16). However, when averaging the training blocks (see inset of Figure 5A), we did found significant differences between AV and A groups (t = 2.09, p = 0.03). In intragroup analysis, comparisons between trials with or without visual stimuli returned no significant effects in groups AV (U = 55.0, p = 0.37) or AVi (U = 59.0, p = 0.47).

**Figure 5.**
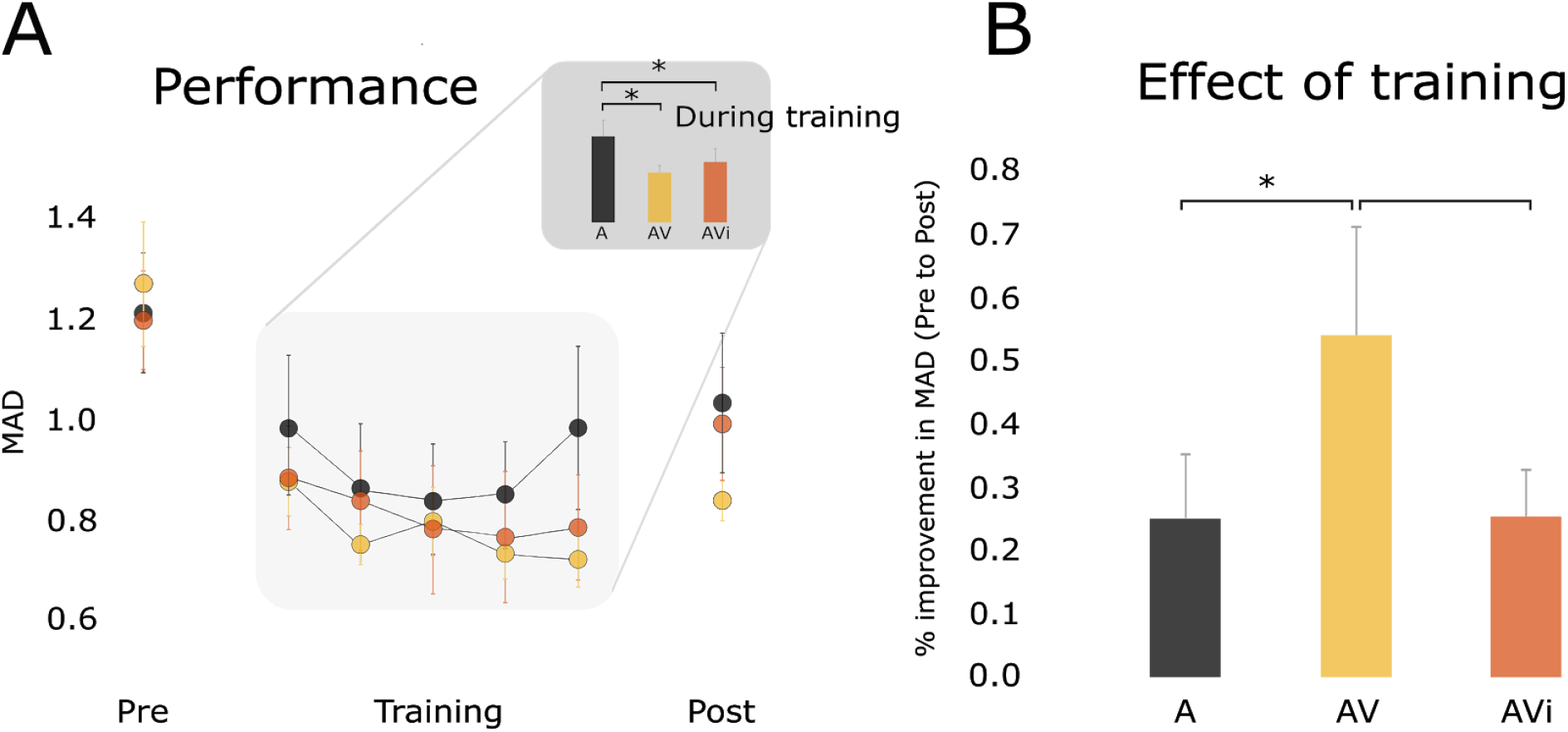
Behavioral performance. **A.** MAD across phases. The inset shows the average from five blocks of training. **B.** Performance improvement (%) from pre-to post-training. *p < 0.05.

In terms of performance improvement from pre- to post-training, we found a significant difference when comparing AV vs. A (Figure 5B) (U = 34.0, p < 0.05). Performance improvement from pre- to post-training was close to significance when comparing groups AV and AVi (U = 37.0, p = 0.06). Finally, groups A and AVi were statistically equal (Figure 5B) (U = 60, p = 0.50).

We also used Cohen’s d as an additional behavioral measurement to compare pre- vs. post-training within and between groups (Table 3). As it can be seen, Cohen’s d support the data shown in Figure 5, revealing very large effect size after training in group AV, whereas small and medium in control groups A an AVi, respectively. Moreover, the comparison between control groups shows a very small effect size, and medium effect size between the test group and both control groups.

**Table 3.** Cohen’s d pre- vs. post-training comparisons

|  |  | Cohen's d | Interpretation |
| --- | --- | --- | --- |
| **Within groups** | $A_{pre} \times A_{post}$ | 0,41 | small |
| | $AV_{pre} \times AV_{post}$ | 1,39 | very large |
| | $AVi_{pre} \times AVi_{post}$ | 0,57 | medium |
| **Between groups** | AV x A | 0,62 | medium |
|  | AV x AVi | 0,60 | medium |
|  | A x AVi | 0,008 | very small |

To further our analysis, we broke down the performance into the pitches. As shown in Figure 6A, MAD from random performance was characterized by a U-shaped quadratic fit across pitches. Pre-training performance also followed a U-shaped curve, whereas post-training MAD followed an inverted U-shaped curve. In an attempt to capture the strongest MAD effects revealed by these fits, we sorted inner (2 to 5) and outer pitches (1 and 6), and then made the group comparisons shown in Figure 6B. For inner pitches, pre- vs. post-training MAD differences were clearly not significant. On the other hand, MAD improvements for outer pitches were significant when comparing A vs. AV (U = 30.0, p = 0.02), and close-to-significant when comparing AV vs. AVi (U = 37.0, p = 0.06).

**Figure 6.**
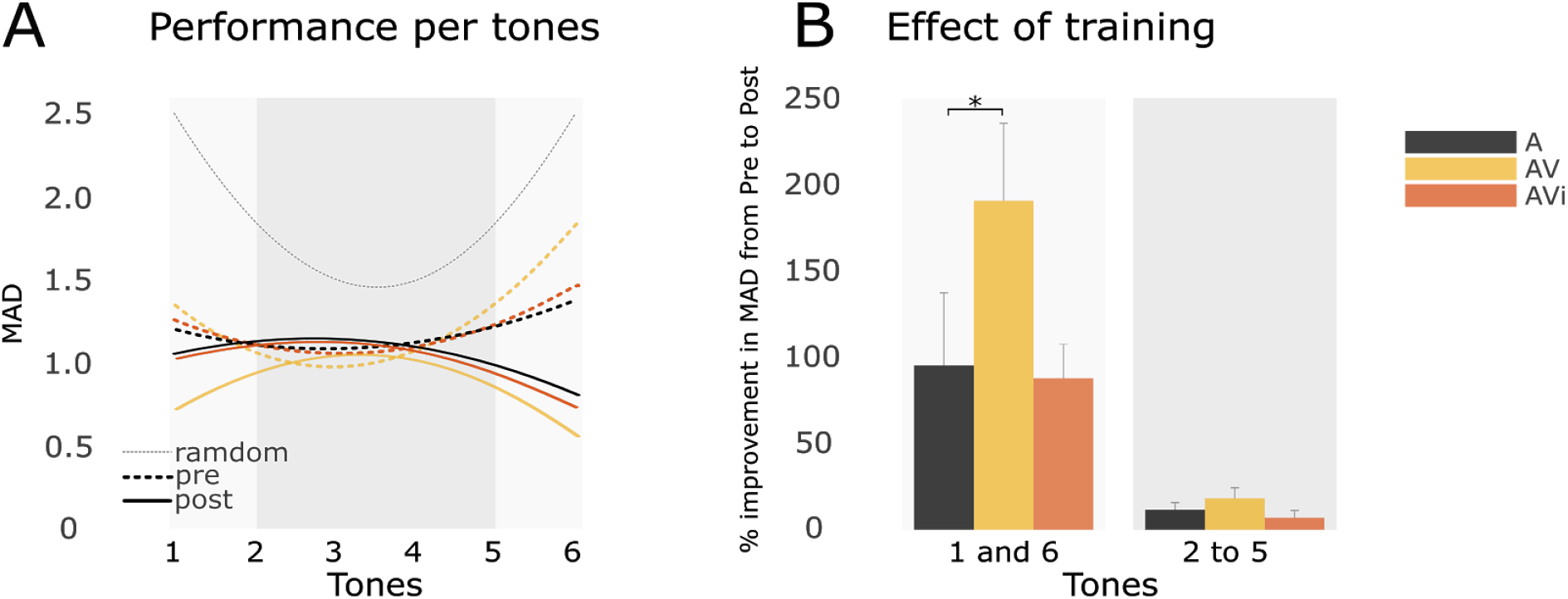
Performances across pitches. **A.** Quadratic fits comparing random (solid gray curve), pre-training (colored dashed curves), and post-training MAD (colored solid curves). Light and dark gray backgrounds correspond to outer and inner pitches, respectively. **B.** Pre-vs. post-training improvements of each group according to inner and outer pitches. Background gray tones correspond to those of panel A. *p < 0.05.

RT did not differ between groups in pre-training, but did differ in the post-training phase (H = 10.79, p < 0,01). We hypothesized this increased response efficacy to be commensurate with the level of confidence. Thus, we compared confidence ratings between groups, but found no effects in either pre- (H = 3.36, p = 0.18), or post-training (H = 4.628, p = 0.11). On average, all groups reported enhanced confidence (Table 04) (A: U = 24.5, p < 0.01; AV: U = 21.5, p < 0.01; AVn: U = 22, p < 0.01), but the increase from pre- to post-training was not different between groups (H = 2.30, p = 0.31). In addition, we explored the possible relationship between RT and confidence ratings, and only found a close-to-significant negative correlation (R = −0.22, p = 0.07).

**Table 4.** Confidence ratings.

|  | **A** | **AV** | **AVi** |
| --- | --- | --- | --- |
| **pre-training** | 5,4 ± 1,50 | 4,1 ± 0,99 | 5,0 ± 2,42 |
| **post-training** | 6,8 ± 0,91 | 5,9 ± 1,45 | 7,2 ± 1,16 |

One may argue that RT could have influenced behavioral results, favoring the group that took more time to respond. However, we did not find a correlation between RT and the effect of training. Another non-significant correlation was between RT and MSI. Closest to significance was Pre-training vs. musical perception (R = −0.29, p = 0.09).

### 3.2. Electrophysiology

We calculated the power of frequencies from 1 to 50 Hz recorded by the central electrode (Cz). Results show a common pattern in all three groups: increase in theta oscillations (especially within the 500 ms post-stimulus period), and decrease in beta oscillations (with an approximate 250 ms latency from stimuli) (Figure 7).

**Figure 7.**
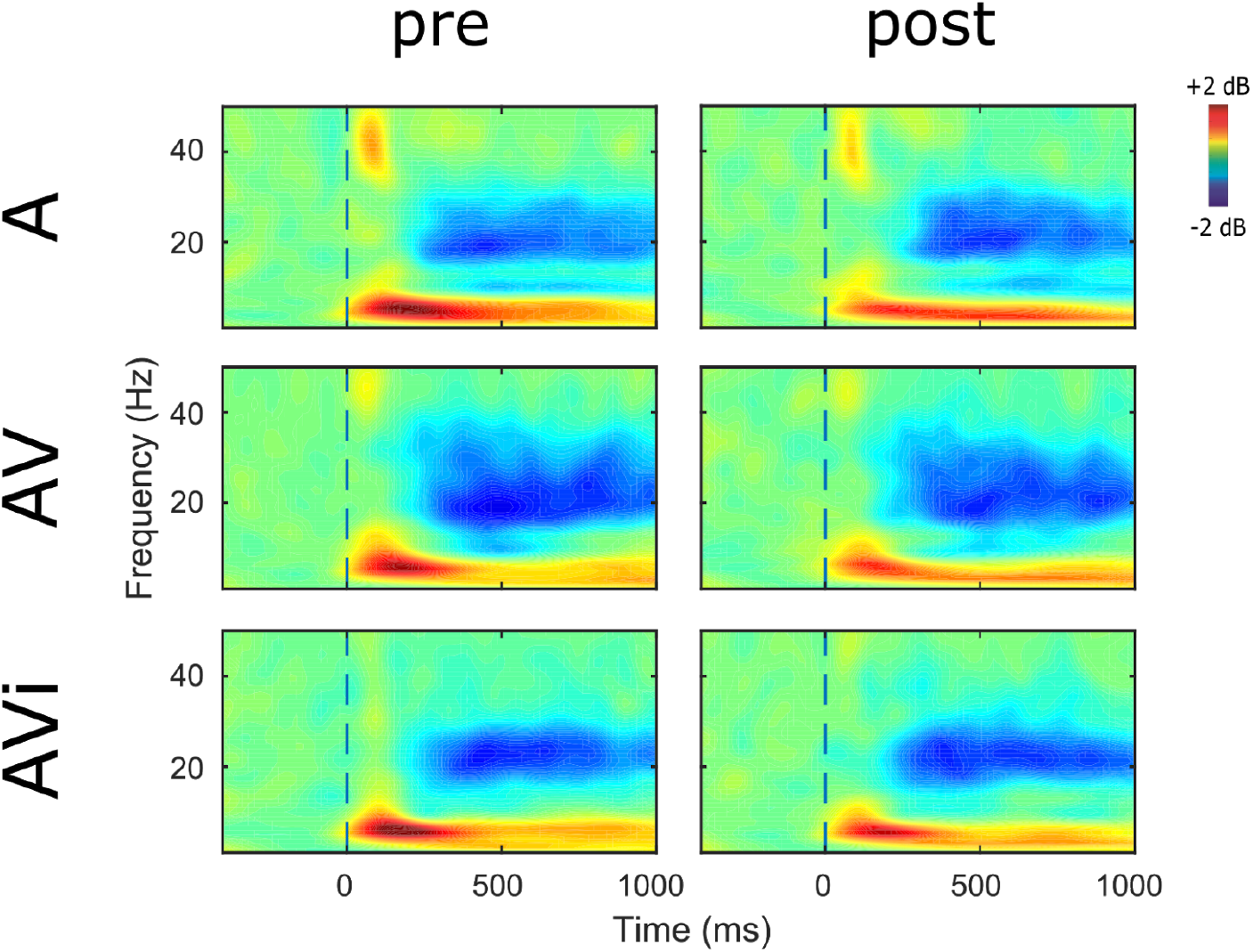
Averaged stimulus-locked spectrograms from the Cz electrode. Left column: pre-training. Right column: post-training. Rows represent the groups. The “zero” time point indicates the stimulus onset. Main effects are seen within the theta (increased power right after stimuli) and beta bands (decreased power from a ~250 ms latency).

After applying the Laplacian transform, we decided to specifically analyze the theta (3-8 Hz), and beta bands (12-30 Hz) by averaging across their frequency bins. We observed the stimulus-evoked reduction of Cz beta power to be statistically equal between pre- and post-training, as well as between groups. Theta analysis, on the other hand, yielded more meaningful findings. As shown in Figure 8, the stimulus-evoked increase in Cz theta power was not uniform across groups. Instead, Group A showed a weaker potentiation of theta, particularly within the 100-250 ms latency from stimuli. This effect was significant when considering all tones (t = 2.42, p < 0.05), or inner tones separately (t = 2.39, p < 0.05). In Group AVi, significant differences were detected only for inner tones (t = 2.60, p < 0.05) (Figure 8).

**Figure 8.**
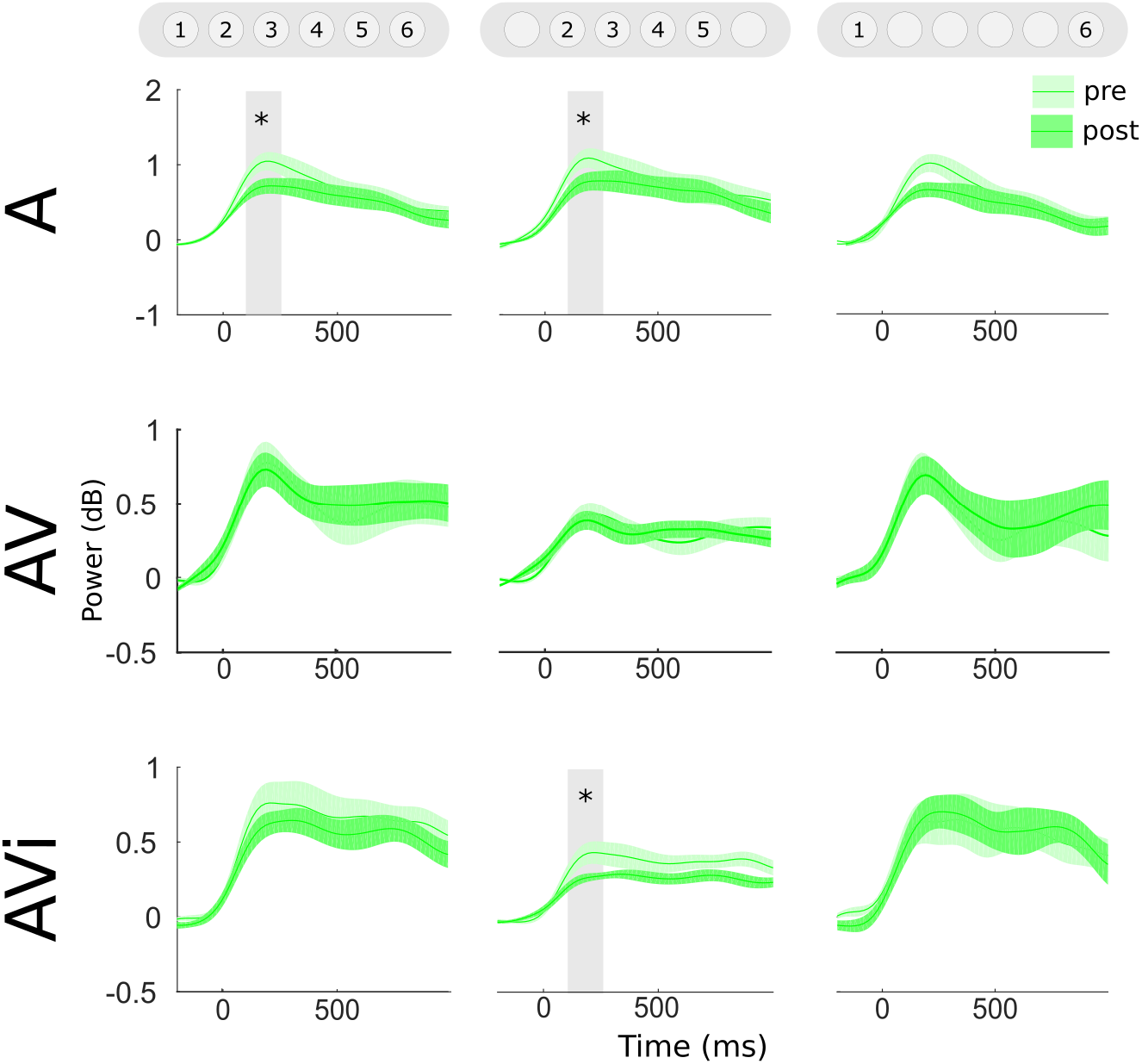
Stimulus-evoked potentiation of Cz theta power (4-8 Hz): comparisons between pre- and post-training. Left column: all tones. Central column: inner tones. Right column: outer tones. Rows represent the groups. Significant differences (p < 0,05) within the 100-250 ms latency are highlighted by gray rectangles.

Between-group comparisons on stimulus-locked band powers can be seen in Figure 9. More specifically, Figure 9 curves are the difference between pre- and post-training data (i.e., post-pre subtraction). The top row of graphs regards 3-5 Hz power in the Cz electrode. Outer pitches evoked an increase that was specific to the AV group and confined to the 200-400 ms latency (F = 3.59, p < 0.05) (Figure 9, right column). The bottom row, in turn, shows 4-8 Hz power in the T7 electrode. Similarly to Cz, we also found an increase specific to the AV group, although at the shorter latency of 100-200 ms (F = 4.16, p < 0.05) (Figure 9, left). Another difference from Cz is that this effect seemed mostly due to inner pitches (F = 3.59, p < 0.05) (Figure 9, center column).

**Figure 9.**
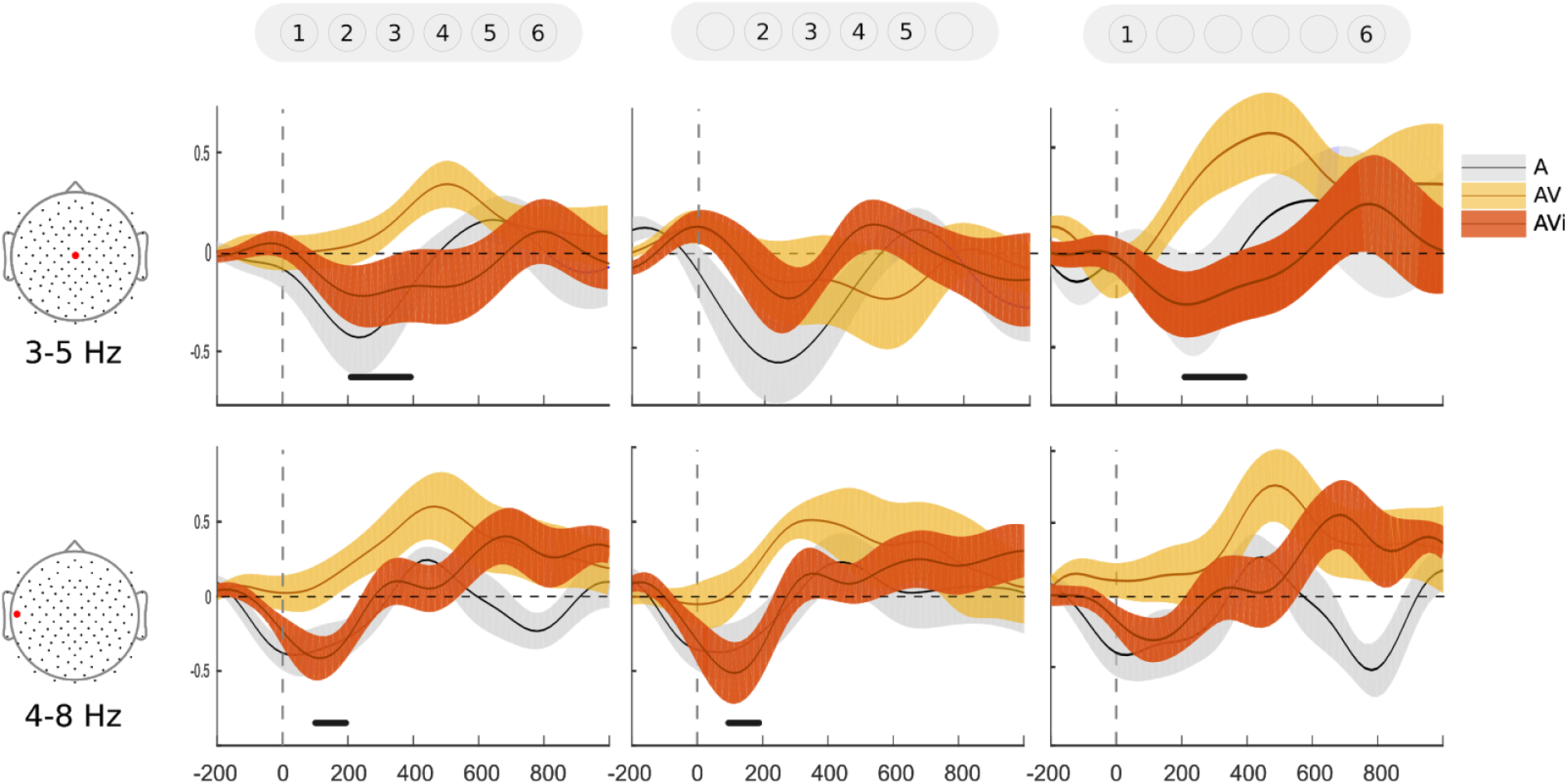
Effects of training on specific bands and electrodes: comparisons between groups. Curves represent the post-pre difference in power. Left column: all pitches. Central column: inner pitches. Right column: outer pitches. Top row: Cz electrode (3-5 Hz). Bottom row: T7 electrode (4-8 Hz). Significant differences are indicated above × axes (p < 0.05).

Lastly, to infer the functional connectivity between auditory and visual cortices, we analyzed the synchrony of representative electrodes placed over the left and right auditory cortices in relation to the whole scalp. Similarly to the analyses above, we also sorted inner and outer pitches. The results show that, in the AV group, theta phase coherence between the left temporal and occipital electrodes increased after training. This effect was observed in a latency window around 140 ms after stimulus onsets (Figure 10). When comparing all groups in this latency window, we observed a marginal effect of group (F = 3.10, p = 0.06). However, when comparing groups in pairs, we found a significant difference in AV vs. A (t = 2.5938, p = 0.0178), and a marginal effect in AV vs. AVi (t = 1.7943, p = 0.0879) (Figure 10A). No significant effects were found in the right hemisphere (Figura 10B), or when separately analyzing the inner pitches.

**Figure 10.**
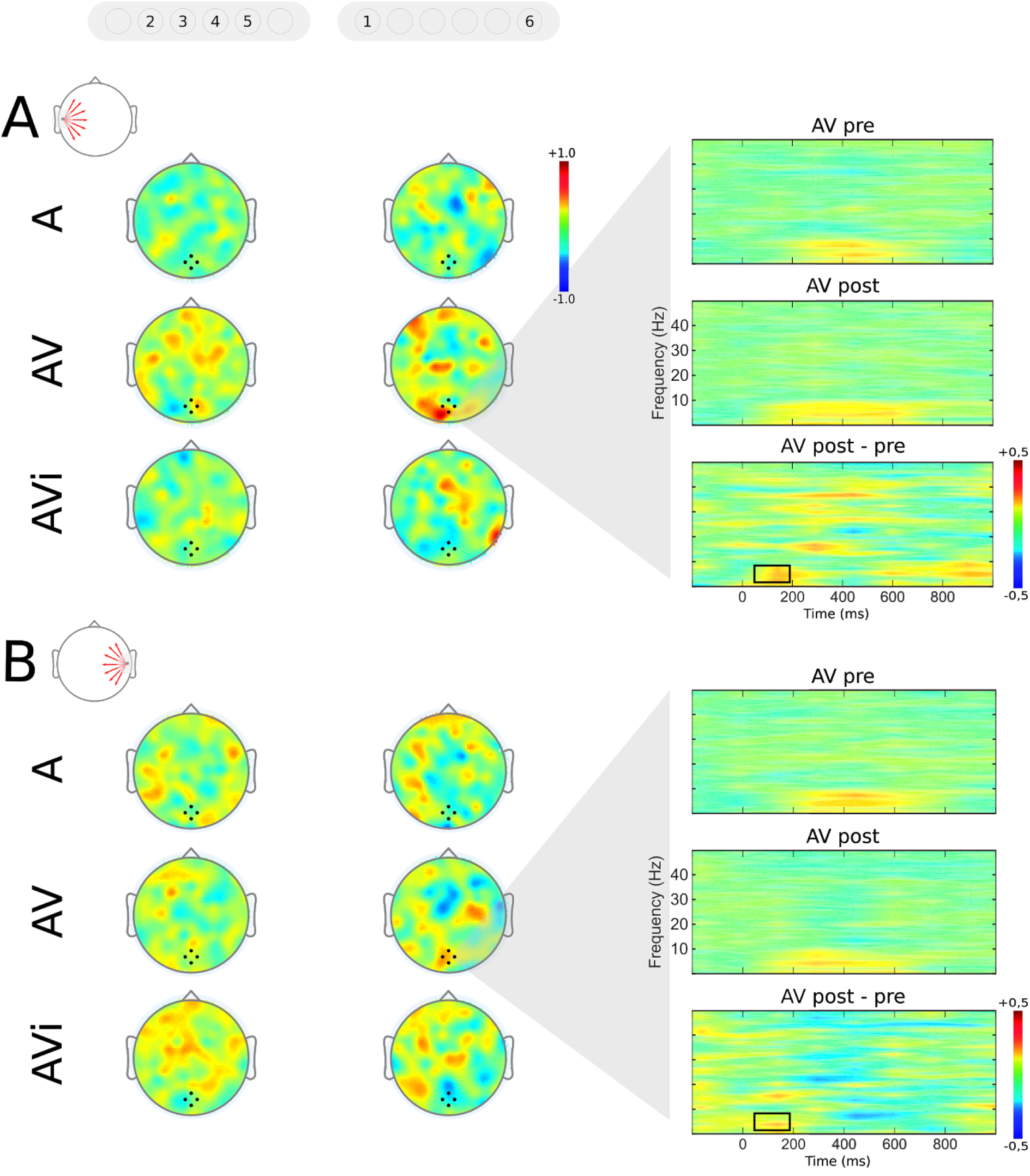
Theta phase coherence between sensory cortices. **A.** Connectivity from T7, over the left temporal cortex. Functional connectivity topoplots were generated for both pre- and post-training (not shown). We then subtracted post-from pre-training to generate the topoplots shown in the figure (i.e., illustrated topoplots represent the post-pre difference in functional connectivity). Topoplot data were averaged from a latency window around 140 ms. Left column of topoplots: inner pitches. Right column: outer pitches. Groups of subjects are indicated from top to bottom. On the right are coherograms from the AV group. From top to bottom, they represent pre-training, post-training, and the post-pre difference, where the ~140 ms effect was found (see black rectangle). **B.** Same as above, but showing connectivity from T8, over the right temporal cortex. No significant effects were found in this case.

## 4. Discussion

In this article we aimed at responding two main questions: 1) Would a pitch learning paradigm benefit from a multisensory training protocol? 2) If so, would early “unisensory” cortices be involved in training-induced alterations in multisensory networks? To approach these questions, we developed a protocol of pitch learning using a short training session (~55 min) with audiovisual stimulation, and with visual stimuli presented below the detection threshold.

### 4.1. Behavioral improvement

The main hypothesis was that the congruent multisensory stimulation would improve perceptual learning, and that early cortices would play a role in this process. Our results show that the test group - trained with auditory plus congruent (and below threshold) visual stimuli - improved better than controls, and further investigation revealed that this improvement was mainly due to responses to the first and last pitches in the herein defined scale, i.e., outer tones. All groups showed improved performance from pre- to post-training phases, and although no significant effects of groups were found, direct comparisons in pairs of groups suggest that the effect of training was stronger in the test group (i.e., with congruent audiovisual stimulation), than in the control groups. From the Cohen’s d perspective (Cohen, 1988; Sawilowsky, 2009), the effect size was considered to be very large in AV and small in both controls (Table 3). Taken together, these two analyses indicate that congruent audiovisual stimulation exerted a distinctive training effect in our experimental design.

Since the mid 2000’s, there has been growing evidence of the influence of multisensory stimulation in learning paradigms manipulating visual (e.g. Seitz et al., 2006, Sham & Seitz, 2008, Deveau et al 2014) and auditory perception (e.g. Lappe et al., 2008; Paraskevopoulos et al., 2012). In general these protocols are conducted for several days, even though it is possible to observe performance differences between uni- and multisensory groups already in the initial sessions (Seitz et al., 2006). Likewise, most researches on auditory learning extend their experiments for several days (e.g. Cuddy, 1970; Menning et al., 2000; Foxton & Brown, 2004; Russo et al., 2005; Seppanen et al., 2012; Paraskevopoulos et al., 2012), although neural plasticity has been documented to take place rapidly, even after a single training session with auditory and visual (Souza et al., 2013) and audiovisual stimulation (Zilber et al., 2014). In this context, it is noteworthy that our results emerged from a very short training protocol: approximately 55 min. This indicates that multisensory perceptual learning in the minutes/hours scale is a promising tool for both academic research and perceptual learning applications.

We observed RT to be different between groups of subjects. The congruent audiovisual group showed similar RT between the pre- and post-training phases, whereas controls responded significantly faster in the post-training. We first hypothesized that confidence ratings could explain this effect, but no significant differences were found. However, there was a trend to lower confidence ratings in the congruent audiovisual group. This could explain higher RT despite the absence of group effects. Interestingly, performance improvements without increases in confidence ratings have been reported in a challenging perceptual task (Zilber et al., 2014). However, the correlation between RT and confidence ratings was negligible in our study. Another hypothesis involving RT is that participants in group AV could have benefited from more time to respond, even though they were instructed to respond as fast as possible. If this was the case, we would expect a positive correlation between RT and the effect of training. However the correlation was also non-significant. The cause of this phenomenon remains open, and further investigation beyond the scope of this paper is needed.

Our behavioral analysis revealed one major source of statistical differences: the performance in trials with outer pitches. In fact, the pair of outer pitches, which accounted for only 33,3% of trials, was the main responsible for pre- to post-training improvements.

Western music is based on scales, in which a group of notes forms a specific structure of intervals. In general, some notes in the scale assume a special status: the tonic note (Herzog, 1949). Still in Western music, the probability of “finding” the tonic note in a melody is higher than that of other notes, and our sensitivity to that probability creates the expectations behind tension and release in melodies (Bigand et al., 1996; Krumhansl et al., 2004). Although the scale we used in this work had no differences in probability, these two pitches (1 and 6) assumed a distinctive status, which possibly created a different expectation compared to the others. Assuming that the perception of a given pitch was influenced by the one presented in the previous trial, one may conclude that any expectation of a tone lower than 1, or higher than 6 was never satisfied. Therefore, both tones shared the fact that any expectation of hearing a tone beyond them in the scale was never satisfied, assuming a different status than the others, like a tonic in a usual Western scale. In sum, we may assume that perceptual learning begins from the outer tones in the scale.

### 4.2. Theta oscillations and early cortices

We found that stimulus-locked theta oscillations varied differently between groups after training. Differences were found between groups in the left temporal electrodes, 100-250 ms after stimulus onsets. Unlike controls, the congruent audiovisual group showed no decrease in theta power from pre- to post-training, and there was a significant difference in inner tones, although with no difference in performance within these pitches. This temporal window, as well as the scalp area where this effect was found, possibly reflects a bottom-up processing. Tesche and Karhu (2000) report that auditory stimulus-evoked theta oscillations were related to neural networks encompassing, for example, the hippocampus. It is indeed expected that tasks like the one used here recruits a large network, consistently with findings that theta oscillations play an integrative role during cognitive demand (Sarnthein et al., 1998; Sauseng et al., 2010; Dai et al., 2017).

In our study, theta oscillations were also significantly different between groups in a later temporal window (200-400 ms), particularly in the test group, central electrodes, and outer tones. These results are inline with the debriefing with participants, that reported being able to identify the outer pitches easily after training. In addition, our findings are consistent with reports on the attention-ERP relationship, given that a well studied ERP component, P300, has a latency that corresponds to the late-onset theta perturbations we observed here (for review, Joos, 2014). Other works show that P300 is related to a higher working memory capacity (e.g., Nittono et al., 1999; Gevins & Smith, 2000), and working memory was, indeed, possibly demanded in the task we used. Some studies support these assumptions: (1) Dong et al. (2015) showed that individuals with higher working memory performance exhibit a greater P300 component, higher theta synchronization, and lower alpha desynchronization (Dong et al., 2015); (2) Gevins and Smith (2000) found a positive correlation between working memory performance and theta synchronization in frontal regions; and (3) Lee et al. (2005) suggested that theta oscillations might be related to maintenance and regulation of relevant information during working memory.

Since theta power in our test group did not decrease from pre- to post-training, this frequency band may have played a role in the performance of these subjects. Another key element corroborating this relationship is that the pre- vs. post-training difference in theta reflects behavioral performance. Specifically in outer-pitch trials, both the auditory-only and audiovisual incongruent groups showed a reduction in theta (although in the audiovisual incongruent group this reduction was weaker), which is in contrast with the effects we observed from the congruent audiovisual group: increased theta power in central, parietal and temporal regions. This scalp distribution suggests that a highly distributed cortical network was involved in this learning protocol, especially in the congruent audiovisual group.

Music and working memory are closely related. Several studies show that musicians perform better than non-musicians in working memory tasks (e.g., Franklin et al., 2008; Parbery-Clark et al., 2009; George & Coch, 2011). However, studies on the neural processes involved in working memory have been controversial. Results from functional MRI and PET studied in the auditory cortex range from activity suppression (Linke et al., 2011; Zatorre et al., 1994) to increase (Linke & Cusack, 2015; Gaab et al., 2003). Recently, Kumar et al. (2016) proposed the separation of encoding, maintenance, and retrieval of information of one among two presented notes. The authors suggest that maintaining a note in working memory involves the auditory cortex, and that sound representations rely on functional connectivities among auditory cortex, hippocampus, and prefrontal cortex (inferior frontal gyrus). Recent studies corroborate this connectivity in its relationship with theta oscillations (Fuentemilla et al., 2014; Herweg et al., 2016; Kaplan et al., 2017; Gruber et al., 2018). Thus, because the task used in our study supposedly demanded maintenance of auditory information, we can propose that the maintenance of theta potentiation in the congruent audiovisual group favored working memory.

Recent studies have shown that the involvement of “unisensory” cortices and their mutual communication have a key role in multisensory integration (Falchier et al., 2002, 2009; Rockland & Ojima, 2003; Cappe & Barone, 2005; Clarke & Innocenti, 1990, Clemo et al., 2008; Vaudano et al., 1991; Laramée et al., 2011, 2013). The multisensory stimulation in our protocol involved nonconscious visual information. By maintaining visual information below detection thresholds we attempted to concentrate visual cortical processing to early stages (Dehaene et al., 2006). If a functional communication exists between early sensory cortices, the activation of visual areas would be enough for the interaction to occur at cortical level. We indeed found evidences of this communication by means of functional connectivity analysis. More specifically, we observed an increase in the post-training connectivity between electrodes over auditory and visual areas, even in phases without visual stimulation. Therefore, we demonstrate that an auditory learning protocol with nonconscious visual stimuli increases synchrony between sensory areas, especially when auditory and visual stimuli are congruent to each other. The fact that this connectivity was increased even in the absence of visual information suggests that plastic changes were stronger, possibly with the recruitment of visual areas for the auditory task after only minutes of training. This corroborates the applicability of multisensory stimulation in short-term perceptual learning protocols.

## 5. Conclusion

The protocol we used for multisensory stimulation favored auditory perceptual learning. Stimulus-locked theta oscillations seemed to play a key role in stimulus processing, which possibly recruited visual areas after training, even in the absence of visual stimuli. In addition, our study may be a first step toward the development of more efficient auditory learning protocols, with eyes open.

